# Prompt-to-Pill: Multi-Agent Drug Discovery and Clinical Simulation Pipeline

**DOI:** 10.1101/2025.08.12.669861

**Authors:** Ivana Vichentijevikj, Kostadin Mishev, Monika Simjanoska Misheva

**Affiliations:** iReason LLC, 3rd Macedonian Brigade 37, 1000, Skopje, North Macedonia,, https://www.ireason.mk/; Faculty of Computer Science and Engineering, Ss. Cyril and Methodius University, Rugjer Boshkovikj 16, P.O. 393, 1000, Skopje, North Macedonia

**Keywords:** Large Language Models, LLMs, Drug Discovery DD, Preclinical Phase, Clinical Phase, Multi-Agent Systems MAS, Prompt-to-Pill, ChatMED

## Abstract

This study presents a comprehensive, modular framework for AI-driven drug discovery (DD) and clinical trial simulation, spanning from target identification to virtual patient recruitment. Synthesized from a systematic analysis of 51 LLM-based systems, the proposed *Prompt-to-Pill* ^*^architecture and corresponding implementation leverages a multi-agent system (MAS) divided into DD, preclinical and clinical phases, coordinated by a central *Orchestrator*. Each phase comprises specialized large language model (LLM) for molecular generation, toxicity screening, docking, trial design, and patient matching. To demonstrate the full pipeline in practice, the well-characterized target Dipeptidyl Peptidase 4 (DPP4) was selected as a representative use case. The process begins with generative molecule creation and proceeds through ADMET evaluation, structure-based docking, and lead optimization. Clinical-phase agents then simulate trial generation, patient eligibility screening using EHRs, and predict trial outcomes. By tightly integrating generative, predictive, and retrieval-based LLM components, this architecture bridges drug discovery and preclinical phase with virtual clinical development, offering a demonstration of how LLM-based agents can operationalize the drug development workflow in silico.

## Introduction

The ability of LLMs to learn from massive datasets and adapt to diverse inputs provides unprecedented capabilities that surpass traditional methods. Their use accelerates decision-making and reduces experimental costs across the development pipeline (1), as evidenced by applications ranging from generating novel molecular structures (2; 3) to predicting pharmacokinetics and toxicity (4; 5), simulating clinical trials (6), and optimizing patient-trial matching (7; 8).

The integration of LLMs into drug development pipelines has gained notable traction, especially across preclinical phases.

Gao et al. proposed a domain-guided MAS for reliable drug-target interaction (DTI) prediction, using a debate-based ensemble of LLMs. The framework partitions the DTI task into protein sequence understanding, drug structure analysis, and binding inference, handled by dedicated agents. Evaluation was conducted on the BindingDB dataset, showing improvements in both accuracy and prediction consistency compared to single-LLM baselines. The system integrates *GPT-4o, LLaMA-3*, and *GLM-4-Plus* (9).

Lee et al. developed *CLADD*, a retrieval-augmented MAS addressing multiple DD tasks. CLADD includes specialized teams for molecular annotation, knowledge graph querying, and prediction synthesis. Evaluations spanned property-specific captioning (BBBP, SIDER, ClinTox, BACE), target identification (DrugBank, KIBA), and toxicity classification. All agents were instantiated with *GPT-4o-mini*, showcasing the utility of general-purpose models when combined with structured RAG mechanisms (10).

Song et al. presented *PharmaSwarm*, a hypothesis-driven agent swarm for therapeutic target and compound identification. The architecture orchestrates three specialized agents (*Terrain2Drug, Market2Drug, Paper2Drug*) and a central evaluator, all integrated via a shared memory and tool-augmented validation layer. Case studies included idiopathic pulmonary fibrosis and triple-negative breast cancer, combining omics analysis, literature mining, and market signals. Agents were powered by *GPT-4, Gemini 2*.*5*, and *TxGemma* (11).

Yang et al. proposed *DrugMCTS*, a novel multi-agent drug repurposing system that incorporates Monte Carlo Tree Search (MCTS) with structured agent workflows. Using *Qwen2*.*5-7B-Instruct* for all agents, the system conducts iterative reasoning across molecule retrieval, analysis, filtering, and protein matching. The framework was benchmarked on DrugBank and KIBA, achieving up to 55.34% recall. A case study involving Equol and CXCR3 showed successful prediction of interaction, supported by AutoDock Vina simulations with a binding score of −8.4 kcal/mol (12).

Inoue et al. introduced *DrugAgent*, an explainable multi-agent reasoning system for drug repurposing. Their architecture coordinates agents handling knowledge graph queries, machine learning scoring, and biomedical literature summarization. Evaluation on a kinase inhibitor dataset revealed strong interpretability and modularity. Detailed ablation studies confirmed that each agent contributes distinctly to the performance. The system employed *GPT-4o, o3-mini*, and *GPT-4o-mini*, and the full pipeline is available open-source (13). Among the surveyed systems, only DrugAgent provides a publicly accessible implementation^1^.

None of the described MASs engages with clinical trial simulation, real-world evidence (RWE), or electronic health records (EHRs), thereby limiting their applicability to the preclinical stage of drug development.

This paper introduces *Prompt-to-Pill*, a unified multi-agent framework build on a systematic analysis of 51 LLM-based studies published between 2022 and 2025. The architecture integrates specialized LLM agents for molecule generation, docking, property prediction, trial construction, patient matching, and outcome forecasting through a central Orchestrator. Unlike prior frameworks confined to molecule-level reasoning, *Prompt-to-Pill* establishes an integrated pipeline from molecular ideation to virtual trial execution, demonstrating how modular LLM agents can operate synergistically within a closed-loop DD and development ecosystem. A complete implementation of the pipeline is available at GitHub^2^.

## Methods

The systematic review was conducted in accordance with the Preferred Reporting Items for Systematic Reviews and Meta-Analyses (PRISMA) guidelines. The PRISMA framework was employed to ensure transparency, methodological rigor, and reproducibility in identifying, screening, and synthesizing eligible studies. A structured multi-stage review process was followed, encompassing database search, eligibility screening, full-text assessment, and data extraction. The complete selection workflow is detailed in the accompanying PRISMA flow diagram depicted in Figure 1.

**Fig. 1.**
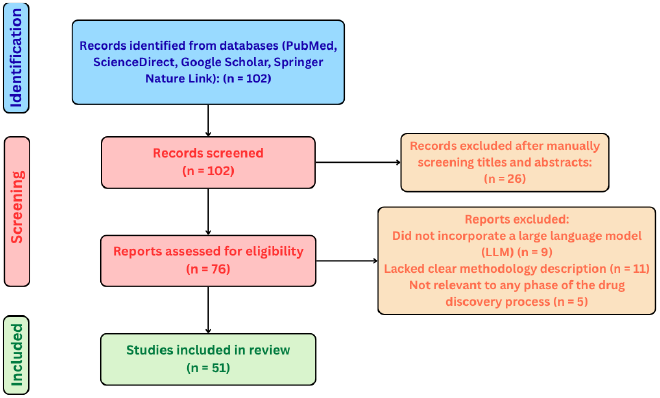
PRISMA-based selection process.

### Information Sources and Search Strategy

A structured and comprehensive literature search was conducted to identify and evaluate LLM-based approaches applied in drug design and discovery. The search was conducted between May 1 and June 15, 2025 across PubMed, ScienceDirect, Google Scholar, and Springer Nature Link. The search covered the publication period 2022 - 2025. Search queries with predefined Boolean combinations captured studies across all DD stages. Representative search strings included: *“large language models” AND (“target identification” OR “binding site prediction”), “large language models” AND (“molecule generation” OR “de novo molecule generation”), “large language models” AND (“clinical trial design” OR “eligibility criteria extraction” OR “trial outcome prediction”), “retrieval-augmented generation” AND (“drug discovery” OR “clinical trials”), “large language models” AND (“patient recruitment” OR “clinical trial matching”* These terms were selected to align with a conceptual pipeline spanning DD, preclinical and clinical phases of pharmaceutical development.

### Study Selection Process

Two reviewers independently screened the titles and abstracts of all retrieved records. Full-text reviews were then performed to assess eligibility based on the predefined inclusion and exclusion criteria.

The inclusion criteria were defined as follows: open-source studies written in English; publications or preprints published between 2022 and 2025; research incorporating LLMs for drug development tasks with clearly defined input–output structure, functional purpose, and workflow integration potential; and studies relevant to at least one stage of the DD or clinical trial process.

The exclusion criteria were: articles not written in English; studies lacking a clear methodological or architectural description; studies that are not publicly accessible; and research not directly applicable to any stage of drug development.

The PRISMA flow diagram in Figure 1 details the number of records identified, screened, excluded (with reasons), and finally taken into consideration for building the Prompt-to-Pill pipeline.

### Data Extraction and Synthesis

For each included study, detailed metadata were manually extracted into structured tables, one for preclinical models and DD and another for clinical applications. Metadata fields were designed to support both technical evaluation and contextual information from each source as follows:

– **Bibliographic**: Authors, Year, Title, DOI.
– **Technical**: Base model (e.g., GPT-4, BioGPT), Task Type, RAG usage, Evaluation Metrics, Datasets.
– **Reproducibility**: GitHub/Hugging Face links, Input/Output examples.
– **Contextual**: Task Narrative, Clinical Trial Phase (I–IV), Abstract Summary.

Extracted data were then synthesized by stage as needed for the drug development pipeline. Studies were profiled and compared across multiple dimensions including application scope, base architecture, task type, and dataset diversity. The The complete metadata tables containing all reviewed studies are provided in the Section 9. This structured comparison informed the construction of the Prompt-to-Pill multi-agent framework introduced later in the paper.

## Methodology

### Prompt-to-Pill Architecture Foundation

The systematic review of 51 studies (2022–2025) shows a sharp growth in research, peaking in 2024 (14 preclinical/DD, 8 clinical), with 17 in 2025, and fewer in 2022 (4) and 2023 (8).

Preclinical/DD studies mainly used generative LLMs such as *LLaMA* and *GPT-2* for creative molecular tasks on open datasets (TDC, DrugBank). Models like *DrugGen* (2) and *DrugGPT* (3) generate SMILES from protein sequences, 3D structures, or text, while others introduce spatial constraints (*3DSMILES-GPT*) or RNA design (*GenerRNA* (15)). *DrugAssist* (16) extends this process with prompt-based molecule optimization, refining compounds to improve pharmacological properties. LLMs also support ADMET prediction, synthesis feasibility, and reactivity analysis (4; 17; 18), as well as biological interaction modeling and drug repurposing (19; 20; 21; 22).

Clinical studies, by contrast, rely on discriminative or hybrid models such as *GPT-4* and *BioBERT*, often trained on structured data (e.g., ClinicalTrials.gov). About half of the 19 clinical papers propose cross-phase models addressing patient selection, outcome prediction, and document generation. LLMs assist in patient-trial matching (7; 8), trial simulation (23; 24; 6), and pharmacovigilance through tools like *AskFDALabel* and *DAEDRA* (25; 26), *occasionally enhanced with RAG pipelines for context-aware text generation (27; 28)*.

Retrieval-augmented generation (RAG) methods showed limited adoption despite their potential for complex reasoning tasks. As shown in Table 1, most studies provided input examples but fewer included output data or reproducible code, underscoring the need for transparency and standardized evaluation.

**Table 1.** Availability of I/O Examples, RAG Usage, and Code Links in Studies.

| Category | Preclinical/DD |  |  | Clinical |  |  |
| --- | --- | --- | --- | --- | --- | --- |
|  | Yes | No | N/A | Yes | No | N/A |
| GitHub/HF link | 20 | 12 | – | 9 | 10 | – |
| Input examples | 17 | 2 | 1 | 6 | 2 | 1 |
| Output examples | 6 | 13 | 1 | 1 | 7 | 1 |
| RAG usage | 3 | 29 | – | 6 | 13 | – |

This analysis informed the design of the proposed Prompt-to-Pill architecture, implemented using the AutoGen (29) framework for scalable multi-agent AI systems. Each agent is adapted from rigorously evaluated domain models, with key performance metrics summarized in Table 2.

**Table 2.**
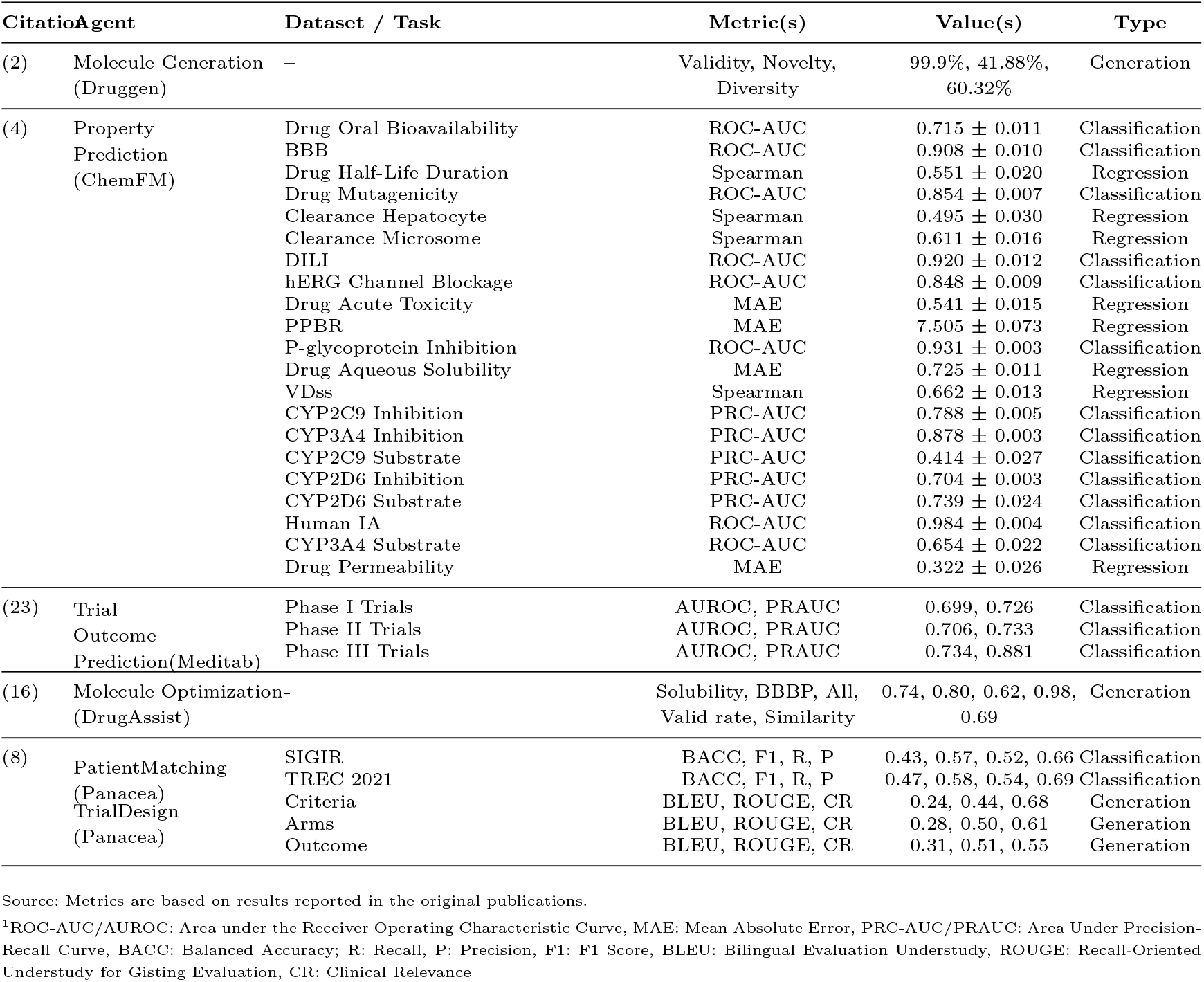
Evaluation metrics of the core agents used in the clinical workflow.

The datasets listed in Table 2 implicitly define the applicability domains (ADs) of the models integrated into the pipeline. For example, ChemFM’s ADME and toxicity predictors are trained on specific benchmarking collections of drug-like compounds, while Panacea and MediTab operate within the disease areas and trial structures represented in CT.gov, SIGIR, and TREC. Because Prompt-to-Pill connects these components sequentially, the effective AD of the full system corresponds to the intersection of all model-specific ADs.

### The Prompt-to-Pill Multi-Agent Pipeline

Constructed from the models identified and reviewed in this study, a comprehensive AI-driven pipeline for DD and clinical trial simulation is presented in Figure 2, structured into three main phases: DD Agents, Preclinical Agents and Clinical Agents, coordinated by a central *Orchestrator*, assisted by a *Planning Agent*. The workflow is task-driven, dynamically selecting the appropriate agent and its tools according to the requirements of the given task.

**Fig. 2.**
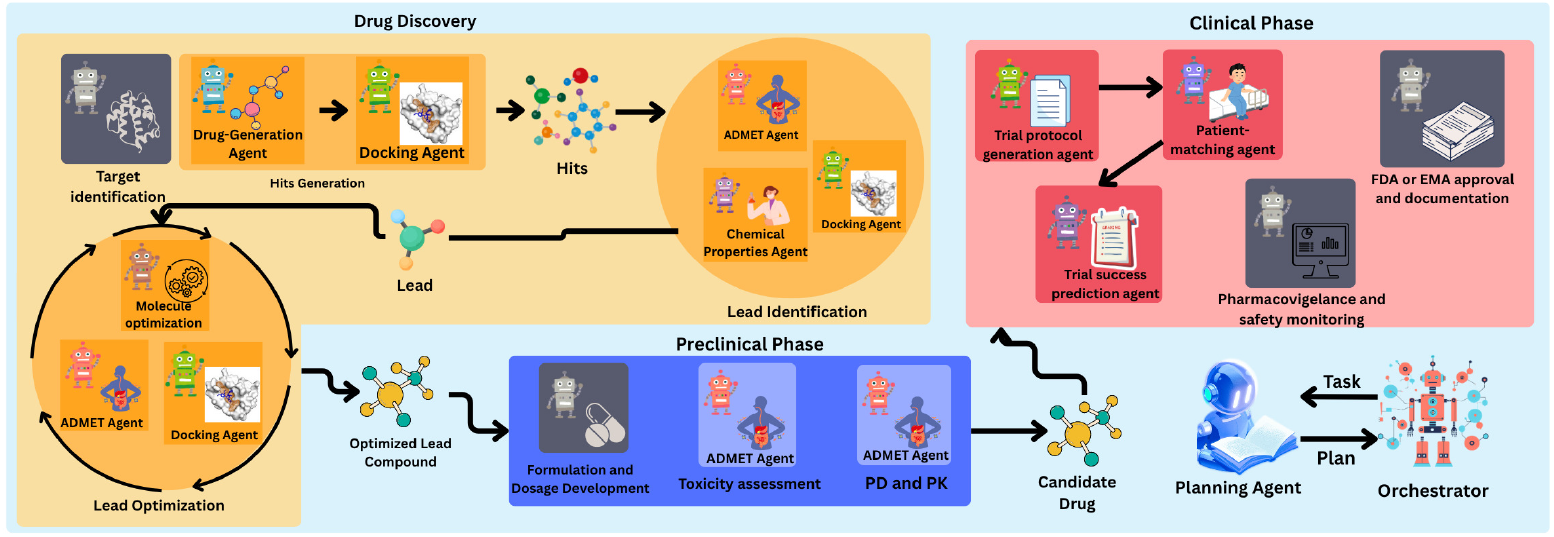
Prompt-to-Pill Multi-Agent Architecture.

In our scenario, we demonstrate this process by focusing on the development of drug candidates for the DPP4 protein target (UniProt ID: P27487). For this drug development task, the pipeline begins with **Drug Discovery Agents**. Here we have 3 subgroups of agents: Hits Generation, Leads Identification and Lead Optimization.

The workflow begins with *Hits Generation*, where the *Drug-Generation Agent*, based on the DrugGen framework (2), produces a set of candidate SMILES sequences. Then the generated SMILES are docked against the target with *Docking Agent*. The Docking Agent is responsible for evaluating the binding affinity of generated molecules against the target protein. Retrieves the target structure from the Protein Data Bank (30) or defaults to AlphaFold models (31) when no experimental structure is available. Candidate SMILES from the *Drug-Generation Agent* are converted into 3D conformations using RDKit^3^. Binding pockets are predicted with P2Rank (32)^4^, and the highest-ranked pocket defines the docking box, whose coordinates are extracted from the P2Rank output and expanded with a fixed padding margin. With receptor and ligand prepared, AutoDock Vina (v1.1.2) performs docking within the predicted pocket, generating 20 poses ranked by affinity. Using this approach, we achieved RMSD lower than 2 Å in 86.59% of cases and a mean RMSD of 1.16 Å on the Astex dataset. The docking setup and visualization, including binding sites, grid box, and ligand, are shown in Figure 3

**Fig. 3.**
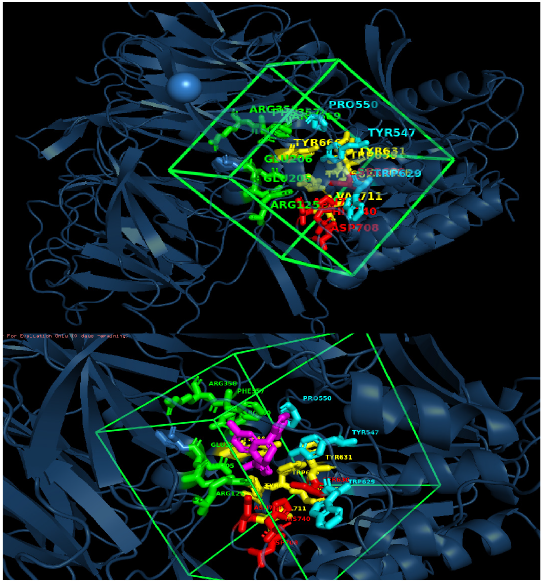
Docking visualization of DPP4 (PDB ID: 2QT9) showing the predicted binding pocket (green grid box; center = 37.87, 49.09, 36.58; edge = 25.02 Å) and the docked ligand (magenta; SMILES: OB(O)c1nnc2n1-c1ccc(Cl)cc1C(c1ccccc1F)=NC2). Pocket side chains are shown as colored sticks (colors for visual separation only) and correspond to validated binding residues ARG125, GLU205, GLU206, TYR547, TYR631, SER630, HIS740, and ASN710 (33).

Following the generation and docking of hits, the workflow progresses to the *Lead Identification* stage. The *Chemical Properties Agent* calculates key physicochemical descriptors (molecular weight, logP, TPSA, hydrogen bond donors and acceptors, rotatable bonds, QED, etc.) using RDkit driven tools. Molecules are filtered according to Lipinski’s Rule of Five^5^ (34) and Veber’s rules^6^ (35), ensuring that only drug-like compounds advance. In parallel, the *ADMET Properties Agent*, using ChemFM(4) framework, is also invoked at this stage to provide an early assessment of ADMET. Properties that this agent can predict are presented in 2. Compound that show the most favorable docking, pass physicochemical filters, and exhibit acceptable ADMET predictions is prioritized as lead.

Next is *Lead Optimizations* stage. This stage focuses on optimizing the chosen molecule to enhance its pharmacological profile while preserving strong binding affinity to the DPP4 target. The *Molecule Optimization Agent*, based on DrugAssist (16), iteratively modifies the structure to enhance bioavailability, solubility, and safety. Each optimized variant is re-evaluated by the *ADMET Properties Agent* and *Docking Agent*, and this loop continues until optimal properties are achieved.

The optimized compound with properties serve entry point into the **Preclinical Phase**, where the optimized candidate undergoes systematic pharmacokinetic and toxicity profiling using ADMET Agent’s tools. Once these evaluations are completed, the workflow is shifted into the **Clinical Phase** for trial simulation.

In the Clinical Phase, the *Trial Generation Agent* constructs a trial protocol tailored to the compound and disease driven by *Panacea* model for criteria, arms and outcomes prediction. This protocol is parsed into structured data and passed to the *Patient-Matching Agent*, which also employs the *Panacea* model (8) to evaluate patient EHR descriptions and identify candidates who meet the trial’s inclusion and exclusion criteria. The agent returns number of matched patients in the final report, and a set of matched patient IDs. These identifiers are saved to a file and the total number of matched patients is computed and included in the final trial report. Subsequently, the *Trial Outcome Prediction Agent* uses *MediTab* (23) to estimate the probability that the proposed trial will succeed, given its protocol structure. In line with the original MediTab formulation(23), this module operates on trial-level metadata and text and learns patterns from historical ClinicalTrials.gov and HINT benchmarks.

Finally, the matched patient data, drug properties, and trial design are provided to the *Orchestrator*, which aggregates all outputs into a structured report.

The input and output format for the Dipeptidyl peptidase 4 (DPP4) target are shown in Figure 4.

**Fig. 4.**
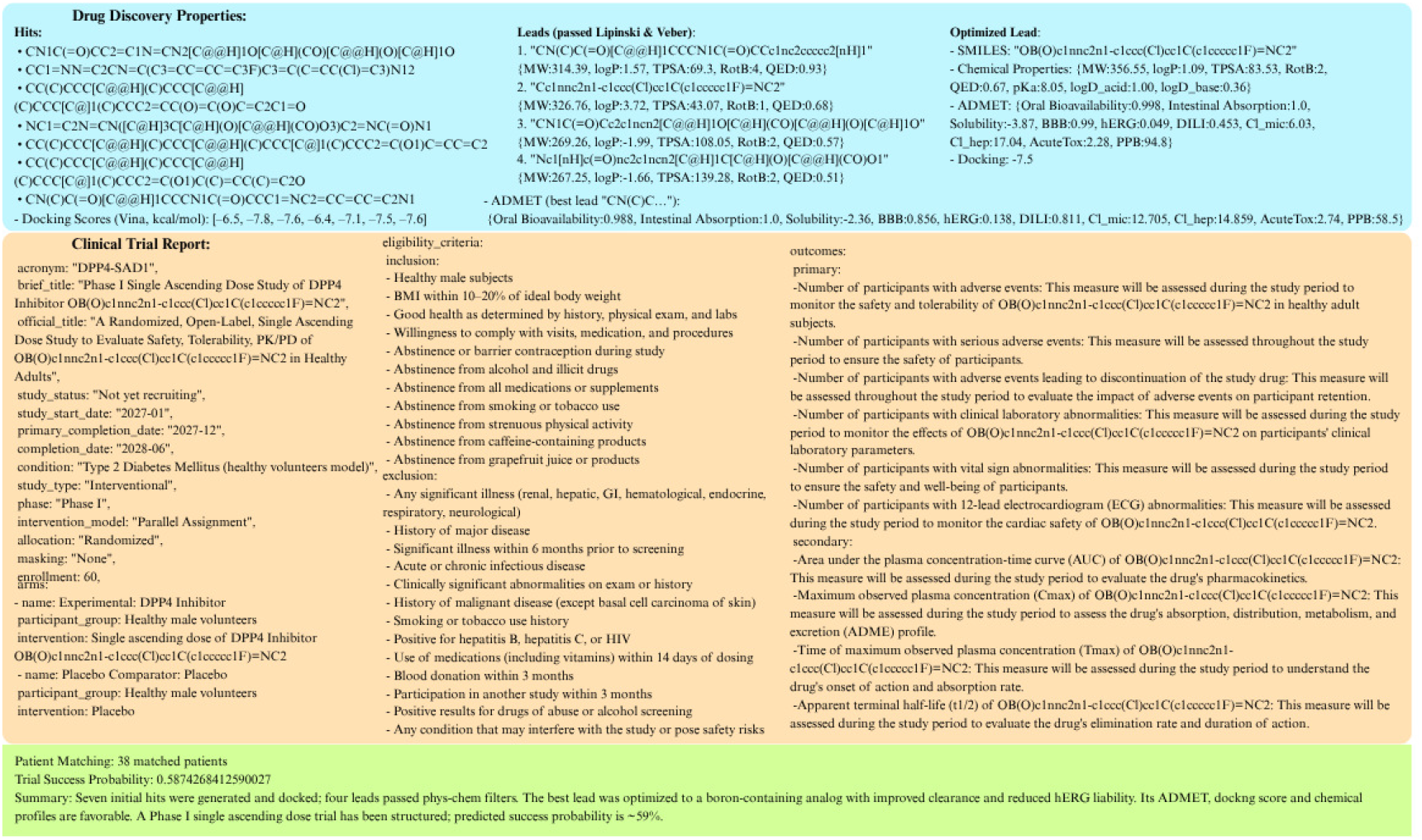
Prompt-to-Pill’s I/O example for Task: “Simulate drug development for DPP4 (P27487) with patients on /path/to/patients.xml”. “Trial Success Probability” correspond to MediTab’s predicted likelihood of clinical trial success based on protocol text and structured trial metadata (e.g., phase, condition, enrollment, arms, outcomes), and do not represent estimates of the underlying drug’s biological efficacy.

## Limitations and Future Work

Prompt-to-Pill is designed as a research-oriented, hypothesis-generation framework intended for use exclusively by trained professionals such as bioinformaticians, medicinal chemists, pharmacologists, and clinicians. The system is not intended for clinical or regulatory decision-making. Instead, its outputs serve as exploratory insights that must be experimentally or clinically validated before any real-world application. The framework aims to support early ideation, academic research, and computational prototyping. While the proposed Prompt-to-Pill pipeline offers a structured, automated approach to drug discovery and clinical simulation, several limitations remain.

First, as shown in Figure 2, some agents remain conceptual placeholders (highlighted in grey), like *Target Identification Agent*, the *Formulation and Dosage Development Agent, FDA or EMA approval and documentation* and the *Pharmacovigilance and Safety Monitoring Agent*. Although we identified related approaches in the literature, most lack accessible implementations or compatible I/O interfaces, preventing integration. Bridging this gap remains a key direction for future work.

Second, the *Orchestrator* is currently implemented using OpenAI’s o4-mini model, which has been shown to perform strongly in medical reasoning and biomedical tasks (36). However, in the trial generation phase, its role is used to producing structured protocol fields such as:study documents, brief summary, acronym, brief title, official title,study status, study start date, primary completion date, completion date, condition, study type, phase, intervention model, allocation, masking, and enrollment. While useful for structuring and simulating trial protocols, these outputs cannot substitute for expert-driven trial design.

Third, each component model operates within the applicability domain (AD) of its training data, and the pipeline therefore inherits the intersection of all ADs. Predictions involving molecules, trial structures, or patient populations far from these distributions should be interpreted as exploratory, not definitive.

This study also presents only a single-target case (DPP-4, P27487), demonstrating feasibility but not generalizability. Future work will extend the framework to multiple targets and diseases for broader validation.

Future work will focus on completing missing agents, enlarging the AD of existing components, and performing multi-disease, multi-target validation studies.

## Discussion

While LLMs have opened transformative opportunities in drug discovery and clinical research, realizing their full potential requires addressing key challenges in transparency, evaluation consistency, and reproducibility.

A key limitation is the limited reasoning ability of current models. In biomedical contexts, correctness alone is insufficient—decisions must be grounded in clear, interpretable reasoning that experts can verify. To address this, several recent models have introduced mechanisms to make reasoning more explicit. These include retrieval-augmented generation (17; 6; 37), instruction-tuned multitask learning (5; 38), and multi-hop rationale generation (37; 39). Such approaches represent important progress toward interpretability. However, without standardized frameworks to assess reasoning quality or consistency, trust in LLM-driven biomedical insights remains limited.

Another major challenge in LLM-based DD is the inconsistency in evaluation protocols across model types. Generative models are assessed using metrics like validity, docking scores, or QED (2; 14; 15), while discriminative models report AUROC or F1 scores (17; 5; 38), yet differ in datasets and thresholds. Knowledge-retrieval and reasoning systems often rely on qualitative outputs without standardized measures (37; 39). This fragmentation hinders comparability and progress. To address this, the field urgently needs task-specific, model-type-sensitive benchmarks.

Reproducibility and transparency also remain persistent issues. Many studies lack public access to code, models, or I/O examples, and when repositories exist, documentation is often incomplete. This fragmentation limits cumulative progress and undermines trust.

These models are the future of drug development, but there is still much work to be done. The path forward requires not only better models, but better systems around the models. This includes standardized evaluations, transparent documentation, expert-guided development, and thoughtful regulation. Only by meeting these unmet needs can we ensure that LLMs evolve from experimental tools to trusted agents in the future of biomedical discovery.

## Ethical and Regulatory Considerations

As highlighted in recent discussions on responsible biomedical AI deployment (40), LLM-based systems in safety-critical domains such as clinical trial design raise central concerns around transparency, explainability, bias mitigation, and the risk of over-reliance on unvalidated outputs. The European Union’s Artificial Intelligence Act (2024) explicitly designates healthcare AI as “high-risk” (Recital 58; Annex III), requiring safeguards such as traceability, human oversight, and fundamental rights impact assessments. The authors have recently published their work on AI Act compliance within the MyHealth@EU framework (41), demonstrating strong ethical responsibility in advancing AI use within sensitive healthcare environments. Their tutorial addresses the dual-compliance challenge of embedding AI Act safeguards (transparency, provenance, robustness) while meeting MyHealth@EU interoperability requirements, showing how AI metadata can be integrated into HL7 CDA and FHIR messages without disrupting existing standards. The goal is not to bypass current guidelines but to ease clinicians’ workload, strengthen trust in AI-assisted decisions, and ensure that compliance and safety are engineered into systems from the outset. In this context, the present pipeline is strictly positioned as a research prototype and decision-support artifact, never as an automated tool for patient eligibility or therapeutic approval. By embedding governance mechanisms early and framing the work as proof-of-concept exploration, the approach contributes to the broader dialogue on trustworthy AI in DD while acknowledging the rigorous benchmarking, reproducibility, and expert oversight still required before clinical translation.

## Conclusion

Prompt-to-Pill introduces a unified multi-agent framework that links the separate stages of drug discovery and clinical development into a coherent workflow. By combining molecular generation, docking, ADMET profiling, trial construction, patient matching, and outcome forecasting within a single orchestrated system, the pipeline demonstrates how diverse LLM-based components can be organized to support early-stage exploration of therapeutic ideas. The DPP4 scenario serves as an example of how a target-specific query can be transformed into a complete in silico development trajectory through modular agent interactions.

As multi-agent methodologies and biomedical language models continue to advance, the future of AI-assisted drug development will depend on interoperable architectures, transparent design principles, and responsible governance. With continued refinement, systems like Prompt-to-Pill may provide a foundation for scalable, hypothesis-driven experimentation across the drug development lifecycle.

## Code Availability

The full implementation of the Prompt-to-Pill multi-agent DD and clinical simulation pipeline is available at the GitHub repository: https://github.com/ChatMED/Prompt-to-Pill.

## Supplementary Material

Table with the retrieved papers for this review is available at the following link: **Supplementary Table 1**.

**Ivana Vichentijevikj** Miss Vichentijevikj is a postgraduate student in Bioinformatics with a biomedical background. She works as a Bioinformatics Software Engineer at iReason, focusing on the development of intelligent systems for drug discovery and clinical data analysis. Her academic and research interests focus on the application of computational methods in drug development and personalized medicine. Ivana is a part of the ChatMED project, where she contributes to the integration of generative AI tools into clinical and pharmaceutical workflows. Her work reflects a strong dedication to advancing translational bioinformatics and promoting the role of AI in the future of healthcare systems.

**Kostadin Mishev**. Prof. Mishev is a professor of Computer Science and Engineering and an expert in the emerging interdisciplinary field that bridges artificial intelligence and DevOps, AIOps, bringing cutting-edge methodologies for operationalizing machine learning pipelines in real-world environments. As a member of the Scientific Committee of the ChatMED project, he contributes strategic and technical expertise in deploying robust, compliant, and scalable AI-driven systems within clinical workflows. Prof. Mishev is also a key industry representative. His portfolio includes the development and deployment of production-grade AI applications, notably, he has led the design of VoiceBot technologies tailored for medical contexts, production-ready for integrating into clinical decision support systems.

**Monika Simjanoska Misheva**. Dr. Simjanoska Misheva is a professor of Computer Science and Engineering with a research focus at the intersection of artificial intelligence, bioinformatics, and clinical decision support systems. She is currently the coordinator of the ChatMED project, a pioneering initiative that aims to integrate generative AI across all levels of a national healthcare system. Her work is driven by the ambition to ensure that the deployment of AI in healthcare is not only innovative but also compliant with the latest European regulations. Dr. Simjanoska’s contributions extend across multi-agent systems, large language models, and their applications in neurology, cancer diagnosis, and personalized medicine. Her recent efforts center on creating regulatory-aware AI architectures capable of multimodal reasoning with genomic, imaging, and biosignal data.

## Footnotes

1 https://anonymous.4open.science/r/DrugAgent-B2EA

2 https://github.com/ChatMED/Prompt-to-Pill

3 https://www.rdkit.org

4 P2Rank success rates: 72.0% Top-n, 78.3% Top-(n+2) on COACH420; 68.6% Top-n, 74.0% Top-(n+2) on HOLO4K

5 HBD ⩽ 5, *HBA* ⩽ 10, *MW* ⩽ 500, *logP* ⩽ 5

6 RotB ⩽ 10, *T PSA* ⩽ 140Å

## References

1. David Oniani, Jordan Hilsman, Chengxi Zang, Junmei Wang, Lianjin Cai, Jan Zawala and Yanshan Wang Emerging Opportunities of Using Large Language Models for Translation Between Drug Molecules and Indications arXiv preprint 2402.09588, 2024.

2. Mahsa Sheikholeslami, Navid Mazrouei, Yousof Gheisari, Afshin Fasihi, Matin Irajpour and Ali Motahharynia. DrugGen enhances drug discovery with large language models and reinforcement learning. Scientific Reports, 15, 13445, 2025.

3. Yuesen Li, Chengyi Gao, Xin Song, Xiangyu Wang, Yungang Xu and Suxia Han. DrugGPT: A GPT-based Strategy for Designing Potential Ligands Targeting Specific Proteins. bioRxiv (Cold Spring Harbor Laboratory), 2023.

4. Feiyang Cai, Katelin Hanna, Tianyu Zhu, Tzuen-Rong Tzeng, Yongping Duan, Ling Liu, Srikanth Pilla, Gang Li, Feng Luo A Foundation Model for Chemical Design and Property Prediction. arXiv preprint 2410.21422, 2025.

5. Yuyan Liu, Sirui Ding, Sheng Zhou, Wenqi Fan and Qiaoyu Tan. MolecularGPT: Open Large Language Model (LLM) for Few-Shot Molecular Property Prediction. arXiv preprint 2406.12950, 2024.

6. Zerui Xu, Fang Wu, Yuanyuan Zhang and Yue Zhao. Retrieval-Reasoning Large Language Model-based Synthetic Clinical Trial Generation. arXiv preprint 2410.12476, 2025.

7. Surabhi Datta, Kyeryoung Lee, Liang-Chin Huang, Hunki Paek, Roger Gildersleeve, Jonathan Gold, Deepak Pillai, Jingqi Wang, Mitchell K. Higashi, Lizheng Shi, Percio S. Gulko, Hua Xu, Chunhua Weng and Xiaoyan Wang. Patient2Trial: From patient to participant in clinical trials using large language models. Informatics in Medicine Unlocked, 101615, 2025.

8. Jiacheng Lin, Hanwen Xu, Zifeng Wang, Sheng Wang and Jimeng Sun. Panacea: A foundation model for clinical trial search, summarization, design, and recruitment. arXiv preprint 2407.11007, 2024.

9. Gao, B., Bai, H., and Zhang, P. [Proposal] A Multi-Agent Framework for Reliable Drug-Target Interaction Prediction. Tsinghua University Course: Advanced Machine Learning.

10. Lee, N., De Brouwer, E., Hajiramezanali, E., Biancalani, T., Park, C., and Scalia, G. (2025). RAG-Enhanced Collaborative LLM Agents for Drug Discovery. preprint, 2502.17506. arXiv

11. Song, K., Trotter, A., and Chen, J. Y. (2025). LLM agent swarm for hypothesis-driven drug discovery. arXiv preprint, 2504.17967.

12. Yang, Z., Wan, Y., Li, Y., Matsuda, Y., Xie, T., and Song, L. (2025). DrugMCTS: A drug repurposing framework combining multi-agent, RAG and Monte Carlo Tree Search. arXiv preprint, 2507.07426.

13. Inoue, Y., Song, T., and Fu, T. (2024). DrugAgent: Explainable drug repurposing agent with large language model-based reasoning. arXiv e-prints, 2408.13378.

14. Jike Wang, Hao Luo, Rui Qin, Mingyang Wang, Xiaozhe Wan, Meijing Fang, Odin Zhang, Qiaolin Gou, Qun Su, Chao Shen, Ziyi You, Liwei Liu, Chang-Yu Hsieh, Tingjun Hou and Yu Kang. 3DSMILES-GPT: 3D molecular pocket-based generation with token-only large language model. Chemical Science, 16:637–648, 2024.

15. Yichong Zhao, Kenta Oono, Hiroki Takizawa and Masaaki Kotera. GenerRNA: A generative pre-trained language model for de novo RNA design. PLoS ONE, 19(10):e0310814, 2024.

16. Geyan Ye, Xibao Cai, Houtim Lai, Xing Wang, Junhong Huang, Longyue Wang, Wei Liu and Xiangxiang Zeng. DrugAssist: a large language model for molecule optimization. Briefings in Bioinformatics, 26, 2024.

17. Eric Wang, Samuel Schmidgall, Paul F. Jaeger, Fan Zhang, Rory Pilgrim, Yossi Matias, Joelle Barral, David Fleet, and Shekoofeh Azizi TxGemma: Efficient and Agentic LLMs for Therapeutics arXiv preprint 2504.06196, 2025.

18. Juan Manuel Zambrano Chaves, Eric Wang, Tao Tu, Eeshit Dhaval Vaishnav, Byron Lee, S. Sara Mahdavi, Christopher Semturs, David Fleet, Vivek Natarajan and Shekoofeh Azizi. Tx-LLM: A Large Language Model for Therapeutics. arXiv preprint 2406.06316, 2024

19. Jon-Michael T. Beasley, Kara Schatz, Elvin Ding, Marcello DeLuca, Nahed Abu Zaid, Nyssa N. Tucker, Rada Y. Chirkova, Daniel J. Crona, Alexander Tropsha and Eugene N. Muratov. TARRAGON: Therapeutic Target Applicability Ranking and Retrieval-Augmented Generation Over Networks./em bioRxiv (Cold Spring Harbor Laboratory), 2025.

20. Ziwen Li, Xiang ‘Anthony’ Chen and Youngseung Jeon. GraPPI: A Retrieve-Divide-Solve GraphRAG Framework for Large-scale Protein-protein Interaction Exploration. arXiv preprint 2501.16382, 2025.

21. Carl Edwards, Aakanksha Naik, Tushar Khot, Martin Burke, Heng Ji, Tom Hope. SynerGPT: In-Context Learning for Personalized Drug Synergy Prediction and Drug Design. bioRxiv (Cold Spring Harbor Laboratory), 2023.

22. Rico Andre Schmitt, Konstantin Buelau, Leon Martin, Christoph Huettl, Michael Schirner, Leon Stefanovski and Petra Ritter Biological Database Mining for LLM-Driven Alzheimer’s Disease Drug Repurposing. bioRxiv (Cold Spring Harbor Laboratory), 2024.

23. Zifeng Wang, Chufan Gao, Cao Xiao and Jimeng Sun. MediTab: Scaling Medical Tabular Data Predictors via Data Consolidation, Enrichment, and Refinement. arXiv preprint 2305.12081, 2024.

24. Michael Reinisch, Jianfeng He, Chenxi Liao, Sauleh Ahmad Siddiqui and Bei Xiao. CTP-LLM: Clinical Trial Phase Transition Prediction Using Large Language Models. arXiv preprint 2408.10995, 2024.

25. Leihong Wu, Hong Fang, Yanyan Qu, Joshua Xu and Weida Tong. Leveraging FDA Labeling Documents and Large Language Model to Enhance Annotation, Profiling, and Classification of Drug Adverse Events with AskFDALabel. Drug Safety, 2025.

26. Chris von Csefalvay. DAEDRA: A language model for predicting outcomes in passive pharmacovigilance reporting. arXiv preprint 2402.10951, 2024.

27. Nigel Markey, Ilyass El-Mansouri, Gaetan Rensonnet, Casper van Langen, and Christoph Meier. From RAGs to riches: Utilizing large language models to write documents for clinical trials. Clinical Trials, 2025.

28. Jeffery L Painter, Venkateswara Rao Chalamalasetti, Raymond Kassekert and Andrew Bate. Automating pharmacovigilance evidence generation: using large language models to produce context-aware structured query language JAMIA Open 8, 2024.

29. Wu, Q., Bansal, G., Zhang, J., Wu, Y., Li, B., Zhu, E., Jiang, L., Zhang, X., Zhang, S., Liu, J., et al. (2024). Autogen: Enabling next-gen LLM applications via multi-agent conversations. In Proceedings of the First Conference on Language Modeling.

30. H.M. Berman, J. Westbrook, Z. Feng, G. Gilliland, T.N. Bhat, H. Weissig, I.N. Shindyalov, P.E. Bourne, The Protein Data Bank Nucleic Acids Research, 28:235–242, 2000, 10.1093/nar/28.1.235.

31. Jumper, J. et al. Highly accurate protein structure prediction with AlphaFold. Nature, 596:583–589, 2021.

32. Radoslav Krivák and David Hoksza P2Rank: machine learning based tool for rapid and accurate prediction of ligand binding sites from protein structure J Cheminform, 10, 39, 2019.

33. Mathur, V., Alam, O., Siddiqui, N., Jha, M., Manaithiya, A., Bawa, S., Sharma, N., Alshehri, S., and Alam, P. Insight into Structure Activity Relationship of DPP-4 Inhibitors for Development of Antidiabetic Agents. Molecules, 28(15):5860, 2023. PMC10420935

34. Lipinski, C. A., Lombardo, F., Dominy, B. W. and Feeney, P. J. Experimental and computational approaches to estimate solubility and permeability in drug discovery and development settings. Advanced drug delivery reviews, 46(1-3), 3–26, 2001.

35. Veber, D. F., Johnson, S. R., Cheng, H. Y., Smith, B. R., Ward, K. W. and Kopple, K. D. Molecular properties that influence the oral bioavailability of drug candidates. Journal of medicinal chemistry, 45(12), 2615–2623, 2002.

36. Rahul K. Arora, Jason Wei, Rebecca Soskin Hicks, Preston Bowman, Joaquin Quiñnero-Candela, Foivos Tsimpourlas, Michael Sharman, Meghan Shah, Andrea Vallone, Alex Beutel, Johannes Heidecke, Karan Singhal HealthBench: Evaluating Large Language Models Towards Improved Human Health. 2505.08775, 2025. arXiv preprint

37. Yichun Feng, Jiawei Wang, Ruikun He, Lu Zhou and Yixue Li. A Retrieval-Augmented Knowledge Mining Method with Deep Thinking LLMs for Biomedical Research and Clinical Support. arXiv preprint 2503.23029, 2025.

38. Tengfei Ma, Xuan Lin, Tianle Li, Chaoyi Li, Long Chen, Peng Zhou, Xibao Cai, Xinyu Yang, Daojian Zeng, Dongsheng Cao and Xiangxiang Zeng. Y-Mol: A Multiscale Biomedical Knowledge-Guided Large Language Model for Drug Development. arXiv preprint 2410.11550, 2024.

39. Zifeng Wang, Cao Xiao and Jimeng Sun AutoTrial: Prompting Language Models for Clinical Trial Design arXiv preprint 2305.11366, 2023.

40. Xiangru Tang, Qiao Jin, Kunlun Zhu, Tongxin Yuan, Yichi Zhang, Wangchunshu Zhou, Meng Qu, Yilun Zhao, Jian Tang, Zhuosheng Zhang, Arman Cohan, Dov Greenbaum, Zhiyong Lu and Mark Gerstein. Risks of AI scientists: prioritizing safeguarding over autonomy. Nature Communications, 16, 8317, 2025.

41. Monika Simjanoska Misheva, Dragan Shahpaski, Jovana Dobreva, Djansel Bukovec, Blagojche Gjorgjioski, Marjan Nikolov, Dalibor Frtunikj, Petre Lameski, Azir Aliu, Kostadin Mishev, & Matjaž Gams. (2025). AI Act Compliance within the MyHealth@EU Framework: A Tutorial. Journal of Medical Internet Research, Advance online publication. 10.2196/81184

